# A combined pipeline for quantitative analysis of human brain cytoarchitecture

**DOI:** 10.1101/2020.08.06.219444

**Authors:** Irene Costantini, Giacomo Mazzamuto, Matteo Roffilli, Annunziatina Laurino, Filippo Maria Castelli, Mattia Neri, Giovanni Lughi, Andrea Simonetto, Erica Lazzeri, Luca Pesce, Christophe Destrieux, Ludovico Silvestri, Valerio Conti, Renzo Guerrini, Francesco S. Pavone

**Affiliations:** European Laboratory for Non-Linear Spectroscopy (LENS), University of Florence, Sesto Fiorentino, Italy; National Institute of Optics (INO), National Research Council (CNR), Sesto Fiorentino, Italy; Bioretics srl, Cesena, Italy; Department of Neurofarba Section of Pharmacology and Toxicology, University of Florence, Italy; Department of Physics, University of Florence, Italy; UMR 1253, iBrain, Université de Tours, Inserm, Tours, France; Pediatric Neurology, Neurogenetics and Neurobiology Unit and Laboratories, A. Meyer Children’s Hospital, University of Florence, Florence, Italy

## Abstract

The 3D analysis of the human brain architecture at cellular resolution is still a big challenge. In this work, we propose a pipeline that solves the problem of performing neuronal mapping in large human brain samples at micrometer resolution. First, we introduce the SWITCH/TDE protocol: a robust methodology to clear and label human brain tissue. Then, we implement the 2.5D method based on a Convolutional Neural Network, to automatically detect and segment all neurons. Our method proved to be highly versatile and was applied successfully on specimens from different areas of the cortex originating from different subjects (young, adult and elderly, both healthy and pathological). We quantitatively evaluate the density and, more importantly, the mean volume of the thousands of neurons identified within the specimens. In conclusion, our pipeline makes it possible to study the structural organization of the brain and expands the histopathological studies to the third dimension.

## 1 Introduction

The three-dimensional reconstruction of large volumes of human brain tissue at cellular resolution remains one of the biggest technical challenges of neuroscience. Nowadays, structural analyses are obtained using traditional processes based on 2D evaluation of thin slices, but they still suffer from significant drawbacks. Such limitations are inherent to the bidimensional nature of the classical slide-based preparation, and include: low sensitivity for sparse features, difficult assessment of dimensions, alteration of morphology, visual artifacts (different orientation or distribution), and sampling bias. Despite the substantial advantages prompted by automatic histology instrumentation and serial sectioning [1], lack of three-dimensionality affects the quality of the produced data and reliability of the analysis.

Recent advances in tissue imaging — in terms of optical clearing, fluorescent staining, and microscopy techniques — have paved the way to high-resolution 3D reconstruction of the brain [2]. Indeed, tissue clearing makes antigens and light penetrate deep inside the sample, enabling fluorescence imaging through high-resolution optical techniques. Multiple methods have been developed to achieve sound clearing and homogeneous staining, but only a few of them have been applied to human tissue. Such samples present specific challenges in comparison to animal models: variability of post-mortem fixation conditions, presence of blood inside the vessels, autofluorescence signals coming from lipofuscin-type pigments, and, finally, needs of exogenous labeling [3]. Alteration of antigens, due to fixation and/or long storage, prevent good immunostaining recognition. Normally, diffusion limits the homogeneous penetration of the dye inside the tissue; voluminous macromolecules, like antibodies, can penetrate only a few dozens of microns inside the sample. Among the various techniques that favor diffusion and increase tissue transparency, tissue transformation techniques such as the CLARITY method [4] and its adaptations have had considerable success. However, they also have limitations. Some were developed for application only to specific samples: e.g. pediatric tissue or controlled post mortem fixation conditions [5, 6]. Others demonstrated compatibility with few antibodies and/or can achieve a staining depth of only a few tens of microns and/or are characterized by very long clearing time [7–12]. Recently, Ku et al. [13] introduced ELAST, a technology that transforms tissues into elastic hydrogel allowing the homogeneous staining of 1 cm-thick sections with various antibodies; however, the preparation of the sample requires sophisticated custom-made equipment and long processing time (20 days). Organic-based techniques were adapted to clear and label human brain tissue, but also in this case they need specific sample preparations: fresh-frozen samples [14], fetal brains [15], or in-situ controlled full body perfusion fixation [16]. In conclusion, up to now, we have no flexible strategy for fast clearing of human brain specimens from different ages, formalin fixed for a long time, and compatible with different antibodies labelling.

An additional consideration that needs to be addressed is that the advances in tissue clearing haven not been followed by innovation on large-scale data analysis and management. High-throughput computational approaches are required to scale-up the processing the significant amount of data produced by 3D anatomical reconstructions obtained by the combination of clearing techniques with high-resolution optical methods. Supervised and semi-supervised methods for localization and segmentation of neuron somata have been proposed, based on advanced classical image processing methods [17, 18], Deep Learning (DL) [19] or combinations of DL with classical processing methods, as described in [20], where “semantic deconvolution” based on Convolutional Neural Networks (CNN) is performed in order to enhance the imaged volumes before applying a mean-shift clustering algorithm. However, this approach enables cell counting but not volume assessment. Semi-supervised region growing approaches [20] use three-dimensional image processing algorithms to find the center of the soma and then repeatedly grow the volume to determine the estimated shapes. Computational issues aside, the main drawbacks of this general method are represented by the need for a precise definition of soma centers, the difficulty of finding all of the centers in a large volume and the complexity of correctly limiting the growth process to an optimal contour. Native Machine Learning techniques such as Convolutional Neural Networks [21] and 3D Convolutional Neural Networks [22], on the other hand, are able to better model visual patterns but are demanding both in terms of the required computing power and the extent of the human-annotated ground truth needed for training (which increases exponentially when moving from the 2D to the 3D domain). Semi-supervised 3D CNN methods have been proposed [19] to alleviate the need for extensive volumetric annotation but they require very high computational power capabilities, suffer from limited scalability and are not able to accurately reconstruct soma surfaces.

Considering the difficulties of human tissue labeling, that decrease the quality of the produced images, and the limits of automatic geometric assessment analysis, the possibility of quantitative evaluates neuronal volumes in human brain reconstruction is still absent. Here, we propose a pipeline that faces different challenges of brain mapping: sample preparation and big data analysis. Indeed, to extract quantitative information not only is it important to optimize each single step, but also to devise a synergic pipeline that integrates together all the different aspects. First, we describe a novel flexible methodology, the SWITCH/TDE protocol, to perform reliable clearing and labeling of human brain tissues. Then, we set up a fast and scalable Machine Learning-based strategy, that we refer to as the 2.5D approach, to perform automated three-dimensional neuronal segmentation and to extract quantitative data.

## 2 Results

### 2.1 The SWITCH/TDE clearing and labeling approach

Penetration of macromolecules and light deep inside the sample are critical processes that are necessary to obtain homogeneous staining of the sample and to reach high transparency, which is essential to perform 3D optical reconstruction with fluorescence imaging. In order to obtain a reliable methodology to label and clear human brain samples from different regions, subjects, and fixation conditions, we modified the SWITCH tissue transformation protocol [9] and we combined it with the 2, 2’-thiodiethanol (TDE) clearing method [5] (Figure 1a). Amongst the various techniques, we decided to use the SWITCH methodology since it allows to control the chemical interaction time and kinetics taking place inside the tissue. By modifying the solutions used during the fixation and clearing, we achieved a more uniform processing of tissues up to 1 mm. At first, we optimized the fixation condition during the SWITCH protocol lowering the concentration of glutaraldehyde (from 1% to 0.5%) during the SWITCH ON step (data not shown). Then, depending on tissue characteristics, we incubated the different samples in the SWITCH clearing solution at 70 °C from 6 hours to one day. Finally, we used the aqueous agent TDE to reduce the Refractive Index (RI) inhomogeneity between the tissue and the surrounding medium, thus minimizing the scattering of light and guaranteeing final transparency of the sample (Figure 1b). Differently from our previous paper [5], we used one-day serial incubations at Room Temperature (RT) up to a concentration of 68% TDE in Phosphate Buffered Saline (PBS) to obtain homogeneous clearing of both grey and white matter. The final solution is characterized by a refractive index of 1.46 equal to that of the UV silica glass used for imaging. The combination of the two techniques allows deep tissue imaging with Two-Photon Fluorescence Microscopy (TPFM): small molecules as SYTOX™ Green can be imaged up to 1 mm in depth, while antibodies can homogeneously label 500 μm-thick slices, respectively (Figures 1c, d). Finally, we demonstrated the compatibility of the SWITCH/TDE method with human brain immunostaining using a variety of different antibodies (Table 1), which were able to stain neuronal cells, GABAergic interneurons and interneurons subtypes, neuronal fibers, glial cells and microvasculature. Incubation time and temperature parameters optimizations are described in the supplementary materials (Supplementary Figure 1). Representative images of the different staining are shown in Figure 1e.

**Figure 1:**
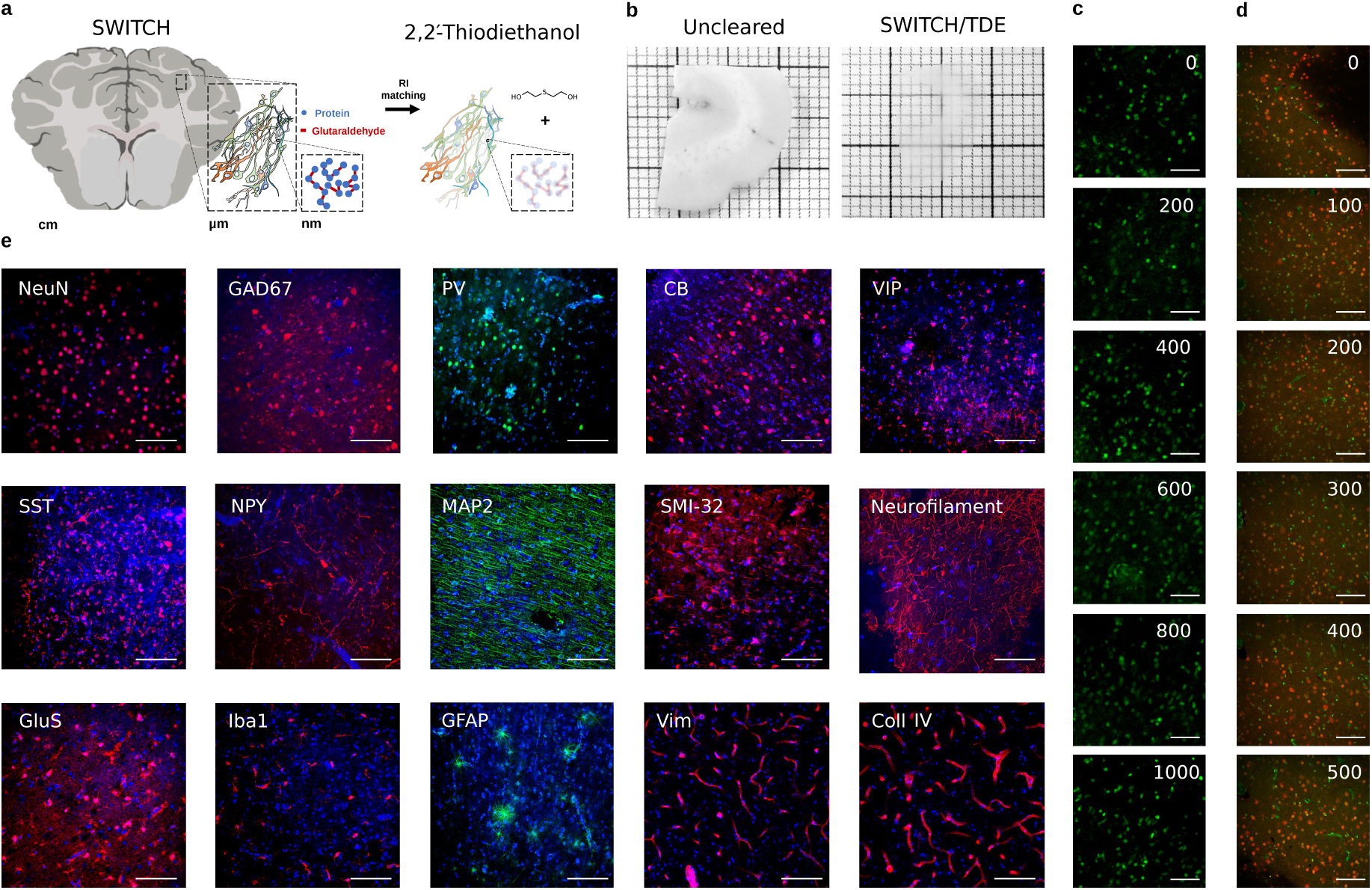
The SWITCH/TDE clearing approach. (a) Schematic illustration of the SWITCH/TDE clearing method. (b) 1 mm-thick slice of an adult human brain sample before and after the treatment. (c) Images of SYTOX™ Green labeled tissue at different depths. Scale bar = 100 μm. (d) Images of NeuN immunostained tissue at different depths. Scale bar = 50 μm. Acquisition obtained with TPFM. (e) Representative images of cleared tissues immunostained with various antibodies and DAPI (4′, 6-Diamidino-2-Phenylindole, Dihydrochloride). Scale bar = 50μm. Acronym list: Neuron-specific Nuclear protein (NeuN, all neurons), Microtubule-Associated Protein 2 (MAP2; pyramidal cells), Nonphosphorylated neurofilament protein (SMI32; pyramidal cells), Glutamic Acid Decarboxylase (GAD67; all GABAergic interneurons), Parvalbumin (PV; GABAergic interneurons subtype), Calbindin (CB; GABAergic interneurons subtype), Vasointestinal peptide (VIP; GABAergic interneurons subtype), Somatostatin (SST; GABAergic interneuron subtype), Neuropeptide Y (NPY; GABAergic interneuron subtype), Neurofilament (NF), Ionized calcium Binding Adaptor molecule 1 (Iba1; glial cells), Glial Fibrillary Acidic Protein (GFAP; glial cells), Glutamine synthetase (GluS), Vimentin (Vim; Microvasculature), Collagen IV (Coll IV; microvasculature).

**Table 1:** Table summarizing the dyes tested in this study. The P/M column denotes polyclonal vs monoclonal antibodies. The same abbreviations used in Figure 1 are used. AF is a shorthand for Alexa Fluor^®^.

| Molecule | Company | Cat. n. | Host | P/M | Dilution |
| --- | --- | --- | --- | --- | --- |
| NeuN | Merck | ABN91 | Chicken | P | 1:50 |
| GAD67 | Santa Cruz | sc-28376 | Mouse | M | 1:200 |
| GAD65/67 | St John's Lab | STJ93195 | Rabbit | P | 1:200 |
| PV | Abcam | ab11427 | Rabbit | P | 1:200 |
| PV | Abcam | ab32895 | Goat | P | 1:200 |
| CB | Abcam | ab207528 | Rabbit | M | 1:200 |
| VIP | Abcam | ab214244 | Rabbit | M | 1:200 |
| SST | Abcam | ab30788 | Rat | M | 1:200 |
| NPY | Abcam | ab6173 | Sheep | P | 1:200 |
| NPY | Abcam | ab112473 | Mouse | M | 1:200 |
| SMI-32 | Merck | NE1023 | Mouse | M | 1:200 |
| Neurofilament | Abcam | ab4680 | Chicken | P | 1:200 |
| GluS | Merck | MAB302 | Mouse | M | 1:200 |
| MAP2 | Abcam | ab5392 | Chicken | P | 1:200 |
| GFAP | Abcam | ab194324 | Rabbit | M | 1:200 |
| Iba1 | Abcam | ab195031 | Rabbit | M | 1:200 |
| Coll IV | Abcam | ab6586 | Rabbit | P | 1:200 |
| Vim | Abcam | ab8069 | Mouse | M | 1:200 |
| Anti-Rat IgG, AF 568 | Abcam | ab175475 | Donkey | P | 1:200 |
| Anti-Rabbit IgG, AF 568 | Abcam | ab175470 | Donkey | P | 1:200 |
| Anti-Chicken IgY, AF 568 | Abcam | ab175711 | Goat | P | 1:200 |
| Anti-Mouse IgG, AF 568 | Abcam | ab175700 | Donkey | P | 1:200 |
| Anti-Sheep IgG, AF 568 | Abcam | ab175712 | Donkey | P | 1:200 |
| Anti-Rabbit IgG, AF 488 | Abcam | ab150077 | Goat | P | 1:200 |
| Anti-Chicken IgY, AF 488 | Abcam | ab150169 | Goat | P | 1:200 |
| DAPI | Thermo Fisher Scientific | D1306 |  |  | 1:1000 |
| SYTOX<sup>TM</sup> Green | Thermo Fisher Scientific | S7020 |  |  | 1:1000 |

### 2.2 3D reconstruction of cerebral cortex samples

The SWITCH/TDE protocol is able to clear different areas of the human brain cortex from subjects of different ages (pediatric, adult, and elderly), obtained from biopsies collected during epilepsy surgery interventions or autopsy stored up to 7 years in formalin. To demonstrate the versatility of the method, four different human brain specimens, from healthy and diseased patients, were analyzed. Two different portions of the left prefrontal cortex from an adult (Male, sample 1) and an elderly subject (Female, 99 years old, no Alzheimer’s disease but initial cognitive decline, no hypertension; sample 2) One dysplastic brain sample from the left temporo-occipital cortex of a 29-year-old man operated to treat drug resistant epilepsy due to focal cortical dysplasia Type IIa (FCDIIa), and one dysplastic brain sample from the left temporo-parietal cortex of an eight-year-old boy operated to treat drug resistant epilepsy due to hemimegalencephaly (HME), respectively samples 3 and 4. The samples were treated with the SWITCH/TDE clearing method and immunolabeled with the Neuron-specific Nuclear protein (NeuN) to detect neurons and DAPI for nuclear staining (samples area of ≈ (1 × 1)cm^2^ and depth of ≈ 500 μm). Imaging was performed with a custom-made Two-Photon Fluorescence Microscope designed to perform mesoscopic reconstruction with a resolution of (0.88 × 0.88 × 2) μm^3^, see Figure 2. After the acquisition, adjacent stacks were aligned and merged together using a custom-made stitching software called ZetaStitcher [23].

**Figure 2:**
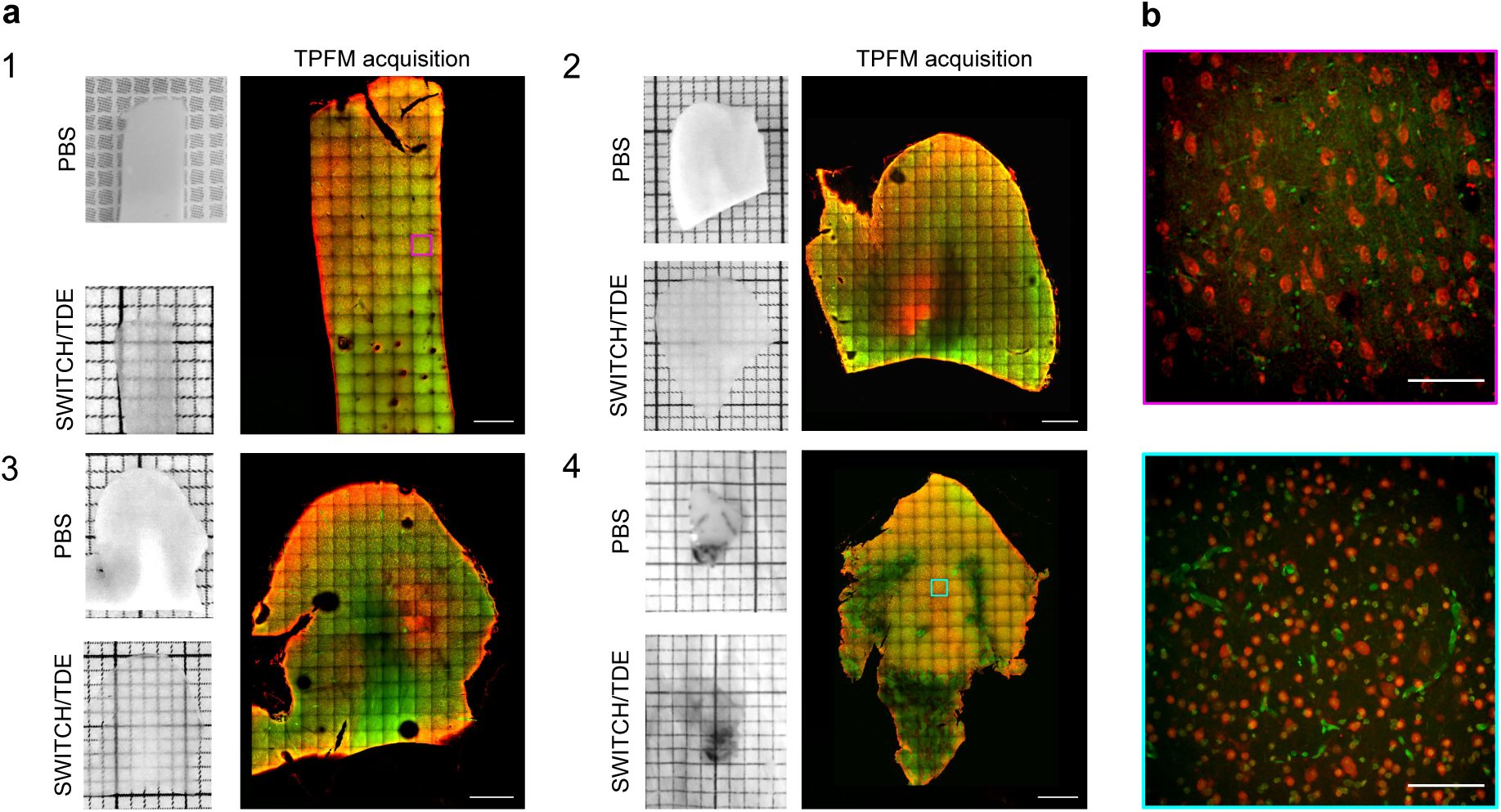
3D mesoscopic reconstruction. (a) Pictures showing the four analyzed human brain specimens before and after SWITCH/TDE clearing. A representative middle plane (*z* ≈ 200 μm) of the mesoscopic reconstruction obtained with TPFM is shown next to each specimen. Scale bar = 1 mm. Specimens 1 and 2: two different portions of the left prefrontal cortex from adult and elderly subjects. Specimens 3 and 4: two surgically removed pieces from patients affected by Focal Cortical Dysplasia Type 2a (FCDIIa) and by Hemimegalencephaly (HME), respectively. (b) Magnified insets of specimen 1 (magenta) and 4 (cyan) showing the native resolution of the acquisition. Tissues were stained with an anti-NeuN antibody (in red) and with DAPI (in green). Scale bar = 100μm

### 2.3 2.5D approach for automatic neuronal volume identification

The volumetric 3D reconstruction obtained with the TPFM consists of tens of GB of data. In particular, the fused volumes of the four samples acquired in this work are sized 19, 50, 57 and 52GB. In order to automatically obtain volumetric information from the 3D reconstruction of the samples imaged with the TPFM, we implemented a novel 2.5D approach based on a Convolutional Neural Network (CNN) for pixel-based classification followed by an analytical reconstruction of 3D polygonal meshes (Figure 3a). The network uses information jointly from the red and green channels (i.e. neurons vs nuclei and tissue autofluorescence) to assign to each pixel a probability of belonging to either the neuron or the background class (Figure 3b and Supplementary video 1). We adopted a pure 2-class fully convolutional CNN that transforms the multichannel source image into a new grayscale one, the so-called probability heatmap. 2D images are processed independently by the neural network, but the resulting heatmaps are reassembled back into a 3D stack, what we refer to as a 2.5D approach. Instance semantic segmentation, based on an iso-surface finding algorithm, is then performed, with a statistical acceptance threshold of 0.5, to the heatmap volume in order to extract the three-dimensional surfaces of each uniquely identified polyhedron.

**Figure 3:**
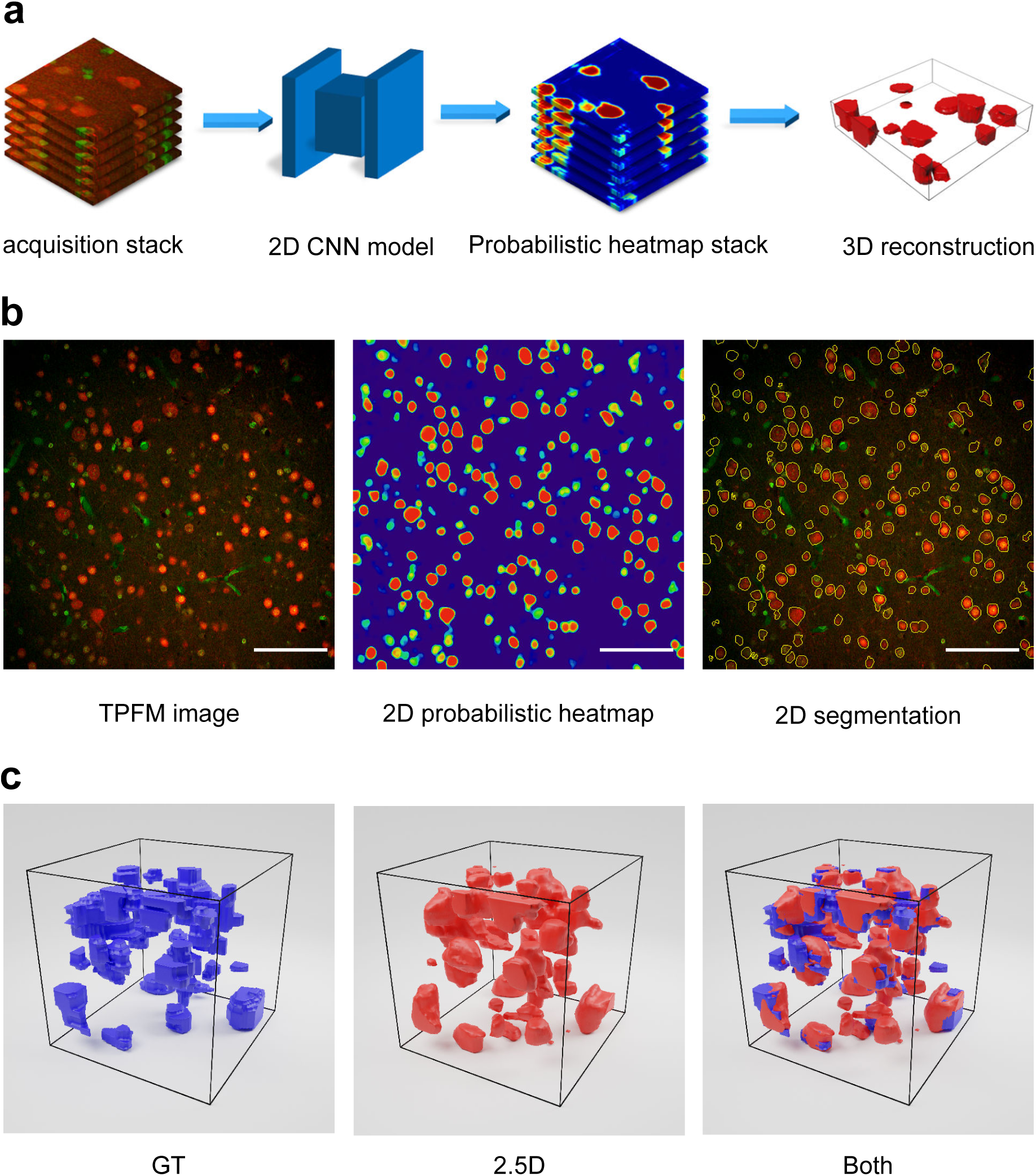
The 2.5D approach. (a) Neuronal segmentation workflow of the 2.5D approach. (b) A representative image undergoing the CNN analysis. From the native image to cells contour segmentation. Scale bar = 100 μm (c) 3D representation of the neurons of a stack manually annotated by the operator (in blue), automatically identified by the 2.5D approach (in red), and the superposition of the two.

The use of a relatively light model (in terms of number of free parameters) allowed us to obtain good segmentation results with advantages on both computational costs and annotation requirements: fast inference times allowed us to obtain results in almost-real-time (with respect to the acquisition time at the microscope) while the number of trainable parameters made it possible to train the model in a supervised fashion using a manageable amount of manually annotated data necessary for the ground truth.

The statistical assessment of the 2.5D performance was determined by analyzing four representative stacks of (100 × 100 × 100) μm^3^, one for each specimen. Each stack was independently manually annotated by an operator and automatically segmented by the 2.5D approach, resulting in a total number of 220 segmented neurons. Figure 3c shows the comparison between the manual annotations and the automatic reconstruction for one of these sub-volumes. For each specimen we first applied a false positive reduction strategy to remove each polyhedron, representing one neuron, with a volume of less than 100 μm^3^, then we calculated the network accuracy on several metrics. The reported values have been computed on a macro statistics basis, i.e. firstly the average of all data of one single specimen is computed and then the average and standard deviation on the four specimens is derived. The volumetric true and false positive fractions designate the extent to which the sub-volume of each neuron detected by the 2.5D segmentation pipeline overlaps with the GT volume or the background and are, respectively equal to (69 ± 6) % and (26 ± 14) %. The total number of neurons found by the 2.5D segmentation is (75 ± 20) % of the true number in the GT. (9 ± 10) % of them can be considered false on macro-average (i.e. they are not present in the GT) while (5 ± 3) % of true neurons are missed (i.e. annotated in the GT but not segmented). Finally, (23 ± 10) % of the identified objects are groups of 2 or more GT neurons merged together into a single object. Indeed, in some circumstances, a single polyhedron found by the network covered more than one real neuron in the ground truth (Supplementary Fig. 3). For each sample, in the supplementary information we report a complete description of all the results.

### 2.4 Neuronal distribution analysis

The four datasets acquired with the TPFM were processed with the 2.5D automatic segmentation method, obtaining meshes for every single neuron in the whole volume (Supplementary video 2). Figure 4 shows a 3D rendering of the meshes for each specimen obtained using the 2.5D approach. The rendering highlights the anatomical architecture of the six cortical layers.

**Figure 4:**
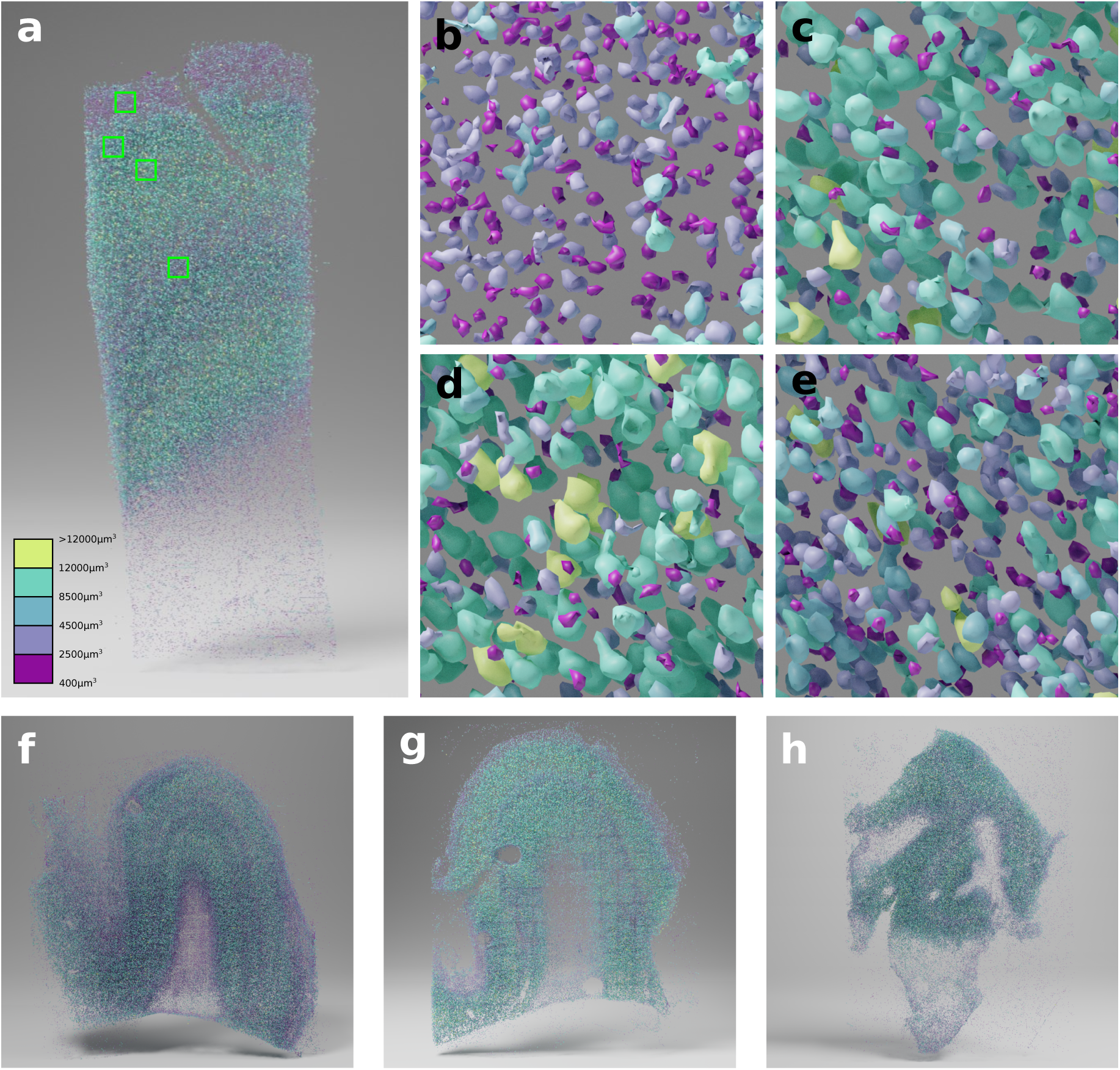
3D rendering of the segmented neurons. Panels a, f, g, h show the 3D rendering for specimens 1, 2, 3 and 4 respectively. The magnified view of the highlighted squares in panel a from top to bottom are shown in panels b, c, d, e, highlighting the neuronal size and density in the different cortical layers.

To quantify the structural organization in the analyzed tissues, we calculated the mean volume distribution and neuronal density distribution. The corresponding maps were obtained, for each specimen, with a binning volume of (100 × 100 × 100) μm^3^ (Figure 5a, b, c). We calculated the densities and the percentage of neuronal volume with respect to the total volume of the grey matter of the sample. To do that, a mask for the grey matter of each samples was manually drawn. To quantify the neuronal distribution along the six cortical layers, masks of each layer volume were manually drawn for each sample. The HME biopsy (sample 4) showed a disruption of the structural organization of the cortex, making layer classification impossible (Supplementary Figure 2). We then measured the volume and density profiles along with cortex depth, which highlight different peaks (Figure 5d). Indeed, volume profiles show peaks in layers 3 and 5, while the neuronal density has a peak in layers 2 and 4. The results of counting are shown in table 2.

**Figure 5:**
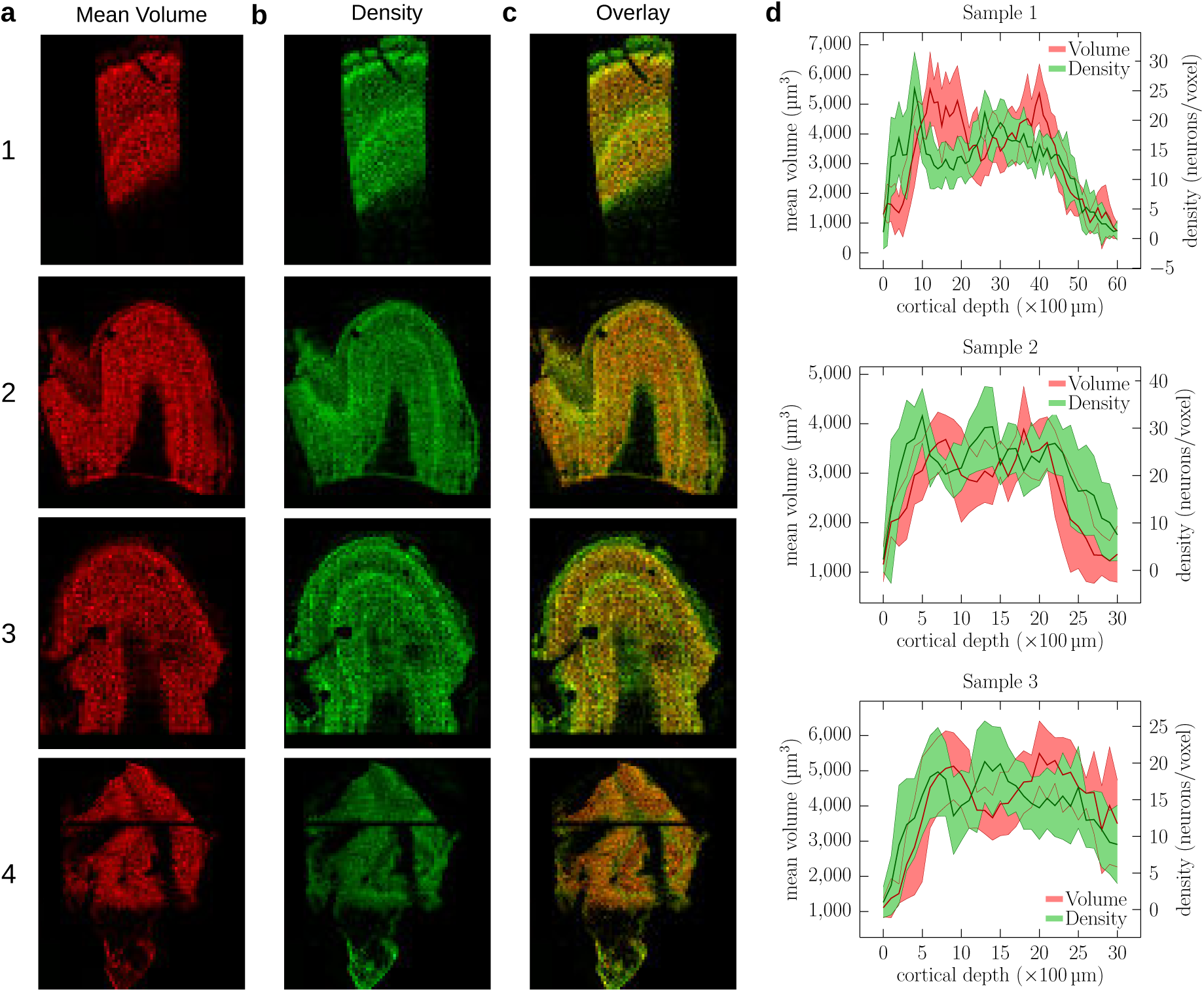
Neuronal distribution analysis. Representative maps of the mean volume (a), neuronal density distribution (b), and overlay of the two maps (c) of the middle plane of each specimen. The maps were computed by performing a 3D binning of 100 μm^3^. Panel (d) shows the neuronal density and mean volume profiles across the cortex as obtained from the maps shown in panels (a) and (b) along 10 different lines that are drawn orthogonal to the cortical layers; the thick line shows the mean value whereas the filled area shows one standard deviation.

**Table 2:** Number, mean volume and density of the neurons in the six layers and in the total volume of the cortex.

|  | Layer | N. Neurons | Tissue volume<br>(mm<sup>3</sup>) | Density<br>(10<sup>3</sup> mm<sup>-3</sup>) | Mean Volume<br>(μm<sup>3</sup>) | Filling<br>fraction |
| --- | --- | --- | --- | --- | --- | --- |
| Specimen 1 |  |  |  |  |  |  |
|  | 1 | 15320 | 2.880 | 5.319 | 2603 | 1.38 % |
|  | 2 | 6454 | 0.486 | 13.280 | 2916 | 3.87 % |
|  | 3 | 19859 | 1.758 | 11.296 | 4441 | 5.02 % |
|  | 4 | 25031 | 1.814 | 13.799 | 3549 | 4.90 % |
|  | 5 | 13972 | 1.300 | 10.748 | 4358 | 4.68 % |
|  | 6 | 6756 | 0.769 | 8.785 | 2263 | 1.99 % |
|  | tot. | 87392 | 9.007 | 9.703 | 3569 | 21.8 % |
| Specimen 2 |  |  |  |  |  |  |
|  | 1 | 69020 | 11.533 | 5.985 | 2550 | 1.5 % |
|  | 2 | 84013 | 11.724 | 7.166 | 2567 | 1.8 % |
|  | 3 | 102169 | 12.789 | 7.989 | 2922 | 2.3 % |
|  | 4 | 57665 | 2.413 | 23.898 | 2730 | 6.5 % |
|  | 5 | 42225 | 2.199 | 19.202 | 3324 | 6.4 % |
|  | 6 | 20289 | 1.054 | 19.250 | 1989 | 3.8 % |
|  | tot. | 375381 | 41.712 | 8.999 | 2740 | 22.4 % |
| Specimen 3 |  |  |  |  |  |  |
|  | 1 | 13815 | 1.646 | 8.393 | 2105 | 1.8 % |
|  | 2 | 22348 | 1.432 | 15.606 | 3412 | 5.3 % |
|  | 3 | 42890 | 3.464 | 12.382 | 4627 | 5.7 % |
|  | 4 | 29834 | 1.804 | 16.538 | 3524 | 5.8 % |
|  | 5 | 33230 | 2.387 | 13.921 | 4584 | 6.4 % |
|  | 6 | 30694 | 5.656 | 5.427 | 3410 | 1.9 % |
|  | tot. | 172811 | 16.389 | 10.544 | 3853 | 26.9 % |
| Specimen 4 | tot. | 177286 | 12.694 | 13.966 | 3008 | 4.2 % |

## 3 Discussion

In this work, we propose a pipeline that addresses some of the most critical challenges of human brain mapping (i.e., sample preparation and big data analysis), enabling a 3D characterization of the cytoarchitecture of the tissue at high resolution. In particular, we develop an approach that allows neuronal segmentation, permitting to evaluate both cell density and mean volumes in mesoscopic reconstruction.

In comparison to animal brains, human neural tissues presents high variability of post-mortem fixation conditions and antigens alterations that prevent proper immunostaining recognition. In this work, we combined the SWITCH tissue transformation method with the TDE clearing. The SWITCH technique allows removing lipids from the sample, while maintaining structural integrity, leading to an increase of tissue permeability and a reduction of the tissue refractive index (RI). The TDE clearing method allows homogenizing the RI of the sample with that of the mounting medium to reach the final transparency. The optimized protocol can perform tissue clearing on prolonged formalin-fixed brain samples and homogeneously stain the tissue with small molecules (up to 1 mm in depth) as well as antibodies (up to 500 μm). The compatibility of the protocol with different antibodies is demonstrated by staining neuronal and non-neuronal cells as well as blood vessels with different antibodies. To illustrate the versatility of the method, we used the SWITCH/TDE approach to prepare volumetric samples (mm^3^-sized volume) from different areas of the cerebral cortex from adult control subjects and pediatric patients with malformations of cortical development. The entire volumes, labeled with anti-NeuN antibody and DAPI, were acquired using a custom-made Two-Photon Fluorescence Microscope (TPFM) capable of performing mesoscopic reconstruction. The optical sectioning and the high-resolution optical investigation made possible by TPFM, in combination with the tissue clearing technique, allowed imaging the 3D organization of whole neurons without introducing any visual artifacts.

Volumetric imaging of samples generates a large amount of data (from tens of GB to tens of TB) that need to be processed in an automated fashion to extract reliable and quantitative information. The software tools we developed in this study made it possible to analyze such big data. As a first step, we stitched together the 3D tiles acquired using the TPFM microscope. Adjacent tiles were aligned and merged by evaluating the cross-correlation of the overlapping areas. Once stitched, we performed an automatic cell segmentation analysis based on a 2.5D Machine Learning approach to achieve a realistic assessment of the neuronal volume. A native 3D implementation of convolutional neural networks, while desirable, is demanding in terms of the required processing power and the extent of the ground truth needed for training. We address the challenges of volumetric segmentation by reformulating the problem as a 2D pixel-based classification task followed by a 3D reconstruction step. The neural network processes each frame independently from the data contained within the previous or the following frame, producing a bidimensional probability map where the value of each pixel is the probability of that very pixel to belong to the foreground class (neuron). By stacking these 2D probability maps, we applied isosurface search algorithms to obtain the 3D representation of the segmented object. While solving a pure 3D problem would imply exploring a cubic space of parameters, our 2.5D reconstruction deals with a quadratic space. Since the number of examples grows exponentially with the space dimensionality, it follows that this 2.5D approach requires much fewer manually annotated examples.

We exploited the 2.5D automatic segmentation method to quantitatively analyze four different specimens (two different samples of prefrontal cortex from an adult and elderly subject, one dysplastic brain sample from the left temporo-occipital cortex of a patient with FCDIIa, and one dysplastic brain sample from the left temporo-parietal cortex of a patient with HME) cleared with the SWITCH/TDE technique and acquired with the TPFM. The 2.5D approach permitted to define the anatomical organization of the neurons in 3D. Indeed, the characterization of density distribution and mean volume allowed us to assess the morphological differences between the arrangement of the layers in the analyzed samples. However, given the small number of samples we analyzed and the general purpose of this study, we did not assess the possible differences between control and dysplastic brain tissues.

In conclusion, we optimized a pipeline that combines the SWITCH/TDE method, a new protocol to clear human brain tissue, with a 2.5D segmentation approach, a technique that makes use of convolutional neural networks to automatically extract information on neuronal volumes and density. The volumetric assessment gives the possibility to extract morphological information that helps discriminating cell types using general staining as NeuN, reducing the labels necessary for the analysis (a critical point in human tissue preparation). Moreover, in the future, the assessment of volume variability could be used in pathological studies to assess, more reliably, the morphological alteration of neurons, increasing the statistical accuracy and the sensitivity of the evaluation. Our work has the purpose of providing a synergic approach enabling a reliable human brain mapping, while addressing the different aspects of quantitative 3D reconstruction analysis. Despite the innovation proposed here, there are still several points that need to be considered to obtain a faster, high-throughput, and informative automated characterization of tissue architecture. A further implementation of the CNN could reduce the errors associated with automatic neuronal counting. At the same time, a combination with a faster optical technique, such as light-sheet microscopy, could facilitate scaling up the analysis. Nevertheless, we believe that our pipeline could be used in the future, not only to provide the anatomical description of samples but also to reduce interpretation biases and to obtain a more precise diagnostic neuropathological assessment.

## 4 Methods

### 4.1 Human brain specimen collection

The study was approved by the Pediatric Ethic Committees of the Tuscany Region (under the project RF-2013-02355240 funded by the Italian Ministry of Health and the Tuscany Region). Healthy tissue samples were obtained from the body donation program (Association des dons du corps) of Universite de Tours and from the Body Donation Program “Donation to Science” of the University of Padova. Prior to death, participants gave their written consent for using their entire body - including the brain - for any educational or research purpose in which anatomy laboratory is involved. The authorization documents (under the form of handwritten testaments) are kept in the files of the Body Donation Program. Pediatric human brain samples were removed during surgical procedures for the treatment of drug-resistant epilepsy in children with malformations of cortical development. Samples were obtained after informed consent, according to the guidelines of the Pediatric Research Ethics Committee of the Tuscany Region. Upon collection, samples were placed in neutral buffered formalin (pH 7.2-7.4) (Diapath, Martinengo, Italy) and stored at room temperature until the transformation and clearing process.

### 4.2 The SWITCH/TDE clearing and labelling protocol

Blocks of fixed samples were washed with a Phosphate Buffered Saline (PBS) solution at 4 °C with gentle shaking for one month to remove formalin from the tissue. Blocks were embedded in a low melting agarose (4% in 0.01 M PBS) and cut into (450 ± 50) μm coronal sections with a vibratome (Vibratome 1000 Plus, Intracel LTD, UK). After the cutting, the agarose surrounding each slice was removed. The permeabilization and staining protocols were modified from that of Murray et al. 2015 [9], as described below. Samples were first incubated in the ice-cold SWITCH-OFF solution (4% GA in PBS 1 X and KHP 0.1 M, titrated with HCl to pH = 3) for 1 day at 4 °C with gentle shaking, then incubated for 1 day in the SWITCH-ON solution (0.5% GA in PBS 1 X, pH = 7.6) for 1 day at 4 °C with gentle shaking. After two washing steps in the PBST solution (PBS with 1% Triton X-100, pH = 7.6) for 4 hours at room temperature (RT), the samples were inactivated with a solution of 4% w/v acetamide and 4% w/v glycine with a pH = 9 (overnight incubation at 37°C). Two washing steps in PBST solution for 4 hours at room temperature (RT) were performed before the incubation in the Clearing Solution (200mM SDS, 20mM Na_2_SO_3_, 20mM H_3_BO_3_, pH = 9) at 70 °C for lipids removal. Incubation time in clearing solution was adapted depending on tissue characteristics: samples from pediatric patients were kept overnight (6-8 hours), while samples from adult and elderly subject up to one day, until complete transparency was achieved. Two washing steps in the PBST solution for 8 hours at room temperature (RT) were performed to prepare the sample for the labeling process. Primary antibodies were incubated in the PBST solution for one day at 4 °C. After two washing steps in the PBST solution for 8 hours at RT, secondary antibodies were incubated in PBST for one day at RT. Table 1 reports the list of antibodies and dilutions used. After two washing steps of 8 hours with PBST at RT, samples were fixed with a 4% solution of paraformaldehyde (PFA) for 10 min at 4 °C to avoid antibody detachment. Samples were then washed three times with PBS for 10 min at RT to remove the excess of PFA. Optical clearing consists in incubation in solutions of increasing concentrations of 20%, 40% and 68% (vol/vol) of 2, 2’-thiodiethanol in 0.01 M PBS (TDE/PBS), each TM for 1 day at room temperature (RT) with gentle shaking. For nuclear staining, DAPI or SYTOX^TM^ Green were diluted in the last incubation of the sample in the 68% (vol/vol) TDE/PBS solution. For 1 mm thick samples, the incubation was performed for two days. Finally, samples were mounted in a custom made chamber with UV silica cover slip (UQG Optics, CFS-5215) that flattens the sample while keeping it completely covered by the TDE/PBS solution allowing a perfect RI matching which is essential for imaging. Sample pictures before and after the clearing process were acquired using a digital camera (Sony DSC-WX500), samples were kept soaked either in PBS or TDE.

### 4.3 Two-Photon Fluorescence Microscopy

A custom-made Two-Photon Fluorescence Microscope (TPFM) was built in order to enable mesoscopic reconstruction of cleared samples. A mode-locked Ti:Sapphire laser (Chameleon, 120 fs pulse width, 80 MHz repetition rate, Coherent, CA) operating at 800 nm was coupled into a custom-made scanning system based on a pair of galvanometric mirrors (LSKGG4/M, Thorlabs, USA). The laser was focused onto the specimen by a refractive index tunable 25× objective lens (LD LCI Plan-Apochromat 25×/0.8 Imm Corr DIC M27, Zeiss, Germany). The system was equipped with a closed-loop XY stage (U-780 PILine^®^ XY Stage System, 135 × 85 mm travel range, Physik Instrumente, Germany) for radial displacement of the sample and with a closed-loop piezoelectric stage (ND72Z2LAQ PIFOC objective scanning system, 2 mm travel range, Physik Instrumente, Germany) for the displacement of the objective along the z-axis. The fluorescence signal was collected by two independent GaAsP photomultiplier modules (H7422, Hamamatsu Photonics, NJ). Emission filters of (440 ± 40) nm, (530 ± 55) nm and (618 ± 25) nm were used to detect the signal, respectively, for DAPI, Sytox Green/Alexa 488, and Alexa Fluor 568. The instrument was controlled by a custom software, written in LabView (National Instruments, TX) able to acquire a whole sample by performing z-stack imaging (depth = (500 ± 100) μm) of adjacent regions with an overlap of 40 μm and a voxel size of (0.88 × 0.88 × 2) μm^3^. The acquisition was performed with a dwell time of 500 μs and the resulting 512 × 512 px images were saved as TIFF files.

### 4.4 Volumetric image stitching

To obtain a single file view of the sample imaged with the TPFM, the acquired stacks were fused together using the ZetaStitcher tool [23]. This software can take advantage of the overlap between neighboring stacks to correct the mechanical error of the imaging platform. Indeed, mesoscopic reconstruction with TPFM can take several days, and temperature changes and mounting medium evaporation can lead to some micron-scale distortion. The software is based on two steps: an alignment process followed by image fusion. As a first step, a 2D cross-correlation map is evaluated at several depths for every pair of adjacent 3D stacks, moving each stack relative to its neighbor. The final position of all stacks is determined by applying a global optimization algorithm to the displacements of the individual pairs. Finally, the stacks are fused into a 3D reconstruction of the whole sample stored in a single TIFF file. The raw datasets of the four samples under investigation in this paper are made available on the EBRAINS platform provided by the Human Brain Project [24].

### 4.5 The Convolutional Neural Network (CNN)

We used a 2D Convolutional Neural Network for pixel-based classification expanding on the design employed in a previous work [25]. In this network architecture, 32 × 32 × 2 sized patches (i.e. considering red and green channels) are extracted from the stitched volume, and fed to the CNN model after a preprocessing step consisting of a single 5 × 5 gaussian kernel filtering stage with *σ* = 3. This operation replicates the intrinsic blurring caused by resampling during data augmentation in the training phase of the CNN. The neural network architecture consists of three convolutional layers, the first two of which are followed by 2 × 2 max-pooling downsampling, and three fully connected layers, the last of which (yellow) makes use of a two-class softmax activation function. A block diagram of the overall network structure is shown in Figure 6. Trainable parameters (205 024 in total) and optimizer hyper-parameters are described in the supplementary information.

**Figure 6:**
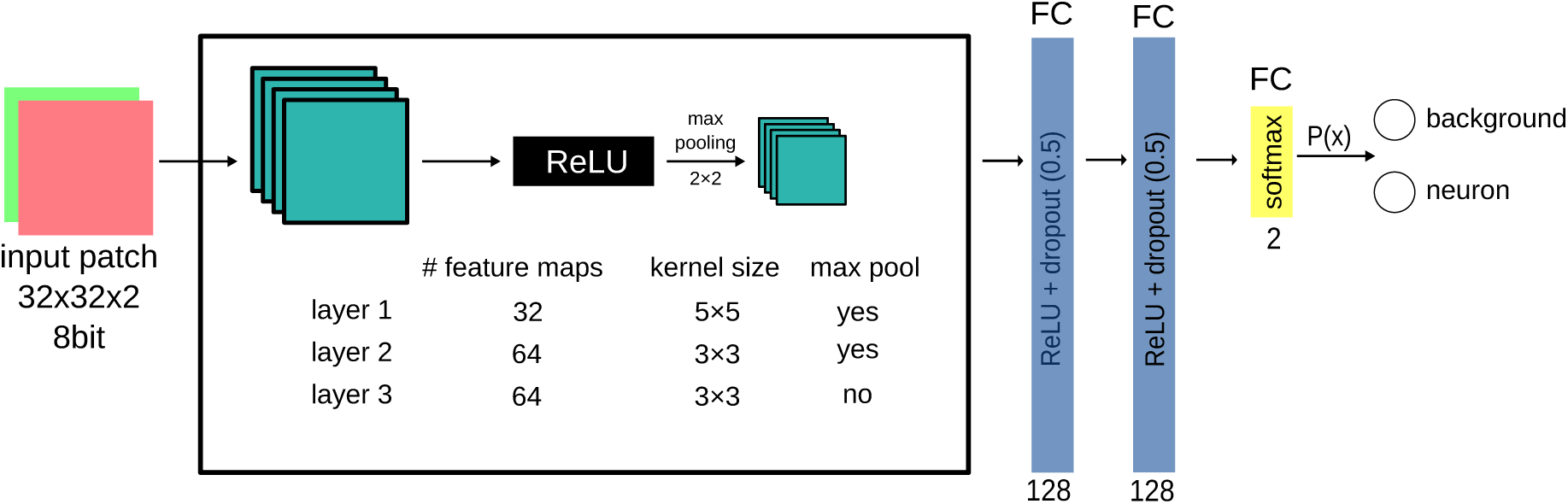
The CNN architecture. Block scheme of the architecture of the CNN with 3 convolutional layers and 3 Fully Connected layers.

The so-defined CNN model classifies the central pixel of each input patch by exploiting the visual pattern of the local neighbourhood (i.e. the coloured 32 × 32 texture) to which the pixel belongs. The model can be used for efficient inference on input data larger than the 32 × 32 patches by exploiting formal equivalence, named fully convolutional, between fully connected layers and 1 × 1 convolutions [26]. This allows us to efficiently produce heatmaps (i.e. probability maps) of entire stack frames.

The ground truth was annotated by two distinct operators on LAIRA^®^ web-based collaborative application [27]. By following an Active Learning paradigm [28] the network was incrementally trained against a number of positive and negative samples originating from the four specimens to improve inter-specimen statistical representatives: the final training dataset is composed of 112 images of 512 × 512px, corresponding to (450 × 450) μm^2^, for a total of 7312 manually annotated neurons (1180 from the first annotation without Active Learning). Additional independent 14 images (1505 neurons) were used to validate the CNN and further 14 images (1208 neurons) to test it. Model regularization is provided in the form of dropout layers, each with a dropout factor of 0.5.

The manually annotated ground truth used to train the neural network is also made available for download on the EBRAINS platform [24] in Ximage format [29] (see Supplementary Information).

### 4.6 The 2.5D approach: from 2D heatmaps to 3D polygon meshes

The CNN model converts entire 2-channel acquisition frames into probability heatmaps; these two-dimensional maps are reassembled back into a 3D stack to obtain an estimate of the three-dimensional probability distribution of neuronal soma presence. The heatmap stack undergoes a post-processing step of false positive reduction represented by the application of a 5 × 5 median filter and a gray-scale morphological opening with a 3 × 3 structuring element. We consider the isosurfaces of this field corresponding to a 0.5 statistical threshold to be representative of the physical boundaries of neuron soma; to calculate them we use a custom variant of the Marching Cubes algorithm [30] followed by additional topological fixes on the identified objects to ensure that every soma is represented by a 2-manifold watertight mesh. This approach allows us to retrieve a three-dimensional vectorial reconstruction of the segmented objects in the entire z-stack, although limited by a grouping effect that sometimes emerges after the instance segmentation step: neurons that are too close to each other are sometimes identified as a single unit (Supplementary Fig. 3). All the 2.5D computations have been performed on a standard Linux-based workstation by the ALIQUIS^®^ software ecosystem [31] with Google TensorFlow as CNN backend [32].

### 4.7 Data analysis

The physical boundaries of the neuronal soma were stored in the form of a triangular meshes in Alembic [33] binary file format, which is suitable for rendering and for further analysis. These files were then processed with Python scripts making use of the trimesh package [34]. The volumes and centroids of all the detected objects were extracted using trimesh.

For neuron counting, volume thresholds were applied to remove segmentation artifacts: volumes lower than 400 μm^3^ and higher than 12000 μm^3^ were discarded. To map neuronal density and volume distribution in the analyzed samples, we plotted 3D histograms with a binning of (100 × 100 × 100) μm^3^ as shown in Figure 5 a, b, c. The centroid value was used to pinpoint the position of the identified neurons within the whole sample volume and in particular to assign each neuron to its corresponding cortical layer according to manually drawn masks. To plot the distributions shown in panel d, 10 different lines were drawn on the binned maps perpendicularly to the cortical layers on which the profiles were extracted. Stacks, 3D stitched volume renderings and videos were obtained using both Fiji [35] and Blender [36].

## Supporting information

Method statistical assessment

## 4.8 Acknowledgments

We thank Raffaele Decaro from the University of Padova for providing a control human brain tissue specimen analyzed in this study. We express our gratitude to the donor involved in the body donation program of the Association des dons du corps du Centre Ouest, Tours, who made this study possible by generously donating his body to science. The research leading to these results has received funding from the European Union’s Horizon 2020 Framework Programme for Research and Innovation under the Specific Grant Agreement No. 785907 (Human Brain Project SGA2) and No. 945539 (Human Brain Project SGA3). This research has also been supported by the Italian Ministry of Health and Tuscany Region (RF-2013-02355240), the Massachusetts General Hospital (The General Hospital Corporation), Athinoula A. Martinos Center, The National Institute of Mental Health (NIMH) through the BRAIN Initiative Cell Census Network under award number 1U01MH117023-01, by the Italian Ministry for Education, University, and Research in the framework of the Eurobioimaging Italian Nodes (ESFRI research infrastructure) - Advanced Light Microscopy Italian Node, and by “Ente Cassa di Risparmio di Firenze” (private foundation) grant n. 24135. Bioretics srl, a company specialized in Machine Learning solutions for Computer Vision, is a subcontractor of LENS in the framework of the BRAIN Initiative Cell Census Network (n. 1U01MH117023-01). The content of this work is solely the responsibility of the authors and does not necessarily represent the official views of the National Institutes of Health.

## 4.9 Author contributions

I.C. and G.M. designed the experiments and made the biological analysis. I.C. developed the SWITCH/TDE method. I.C., E.L., A.L., and L.P. performed sample preparation. I.C. conducted TPFM imaging. G.M. developed the ZetaStitcher software, performed the mesoscopic reconstruction and curated the data. M.R. conceived the 2.5D approach. M.R., G.M., F.C., and I.C. oversaw the overall development of the 2.5D processing pipeline. G.L., and A.S. performed the 3D reconstruction, M.N. operated and tested the neural network. I.C. and A.L. annotated the ground truth, F.C. prepared the 3D renderings. C.D. provided the elderly human brain tissue specimen. V.C. and R.G. provided the pediatric human brain tissue specimens and contributed to the concept of the biological evaluation. I.C., G.M., L.S., and F.S.P. supervised the project. I.C. and G.M. wrote the paper with inputs from all authors.

## 4.11 Data availability

Data supporting the findings of this study are included in figures and videos as representative images or data points in the plots. The raw data of the mesoscopic reconstructions and the ground truth masks are available on the EBRAINS platform at URLs provided in the Methods. Additional images other than the representative images are available from the corresponding author upon reasonable request.

## 4.12 Code availability

ZetaStitcher, the CNN model, and 2.5D approach codes are open source and available at URLs provided in the methods and in the supplementary information. The other custom codes used in this study are available from the corresponding author upon reasonable request.

