## Supplementary Information for "A combined pipeline for quantitative analysis of human brain cytoarchitecture"

#### 1. Immunostaining protocol optimization

Trials with different temperatures (37°C, @RT, and 4°C) and time of incubation (1 or 2 days) were performed in order to identify the best condition for optimal immunostaining. For each condition, images were acquired with the TPFM maintaining the same PMT gain and laser power. Signal to Background (S/B) analysis was performed with Fiji (http://fiji.sc/Fiji) to assess the best contrast: mean intensity of 10 square of 25px were analyzed for each condition, media and standard deviation were then calculated using OriginPro 9.0 (OriginLab Corporation). The protocol with the highest signal amplification corresponds to 4°C of incubation temperature and 24 hours of incubation time of the primary antibody.

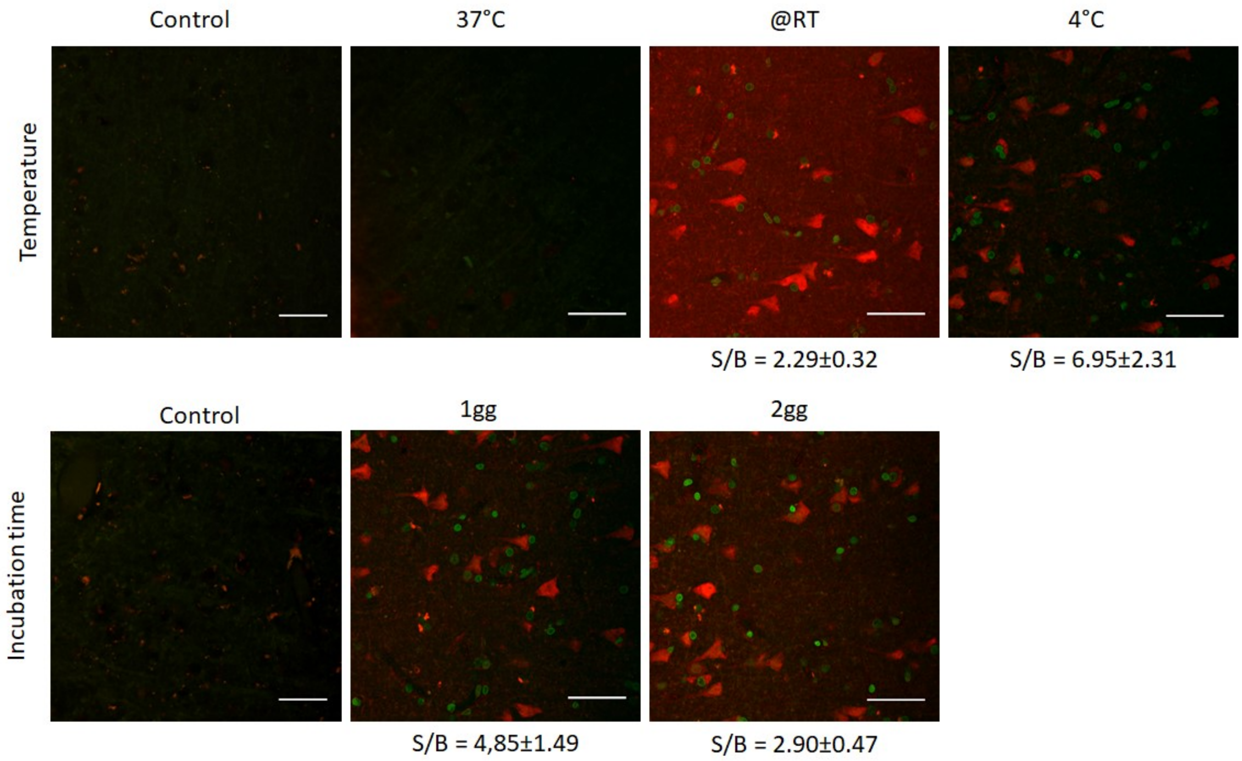


**Supplementary Figure 1**: **Immunostaining optimization**. Representative two-photon images of tissue stained with anti-NeuN antibody (in red) and DAPI (in green) at different temperatures and incubation times. Scale bar = 50 µm.

#### 2. Manual segmentation of the grey matter

In order to identify the different layers of the grey matter of the cortex, the maps obtained analyzing the mean volume distribution and the density distribution with a binning of 100 x 100 x 100 $\mu m^3$ were manually segmented using the software Fiji (Supplementary figure 2). Big blood vessels, tissue holes/breakages, and imaging artifacts were not reckoned drawing the masks. Sample 4 shows a disruption of the structural organization of the cortex, making layer classification impossible, only grey matter were segmented. The total volume for each mask was obtained performing this operation for all the z-plane of the sample.

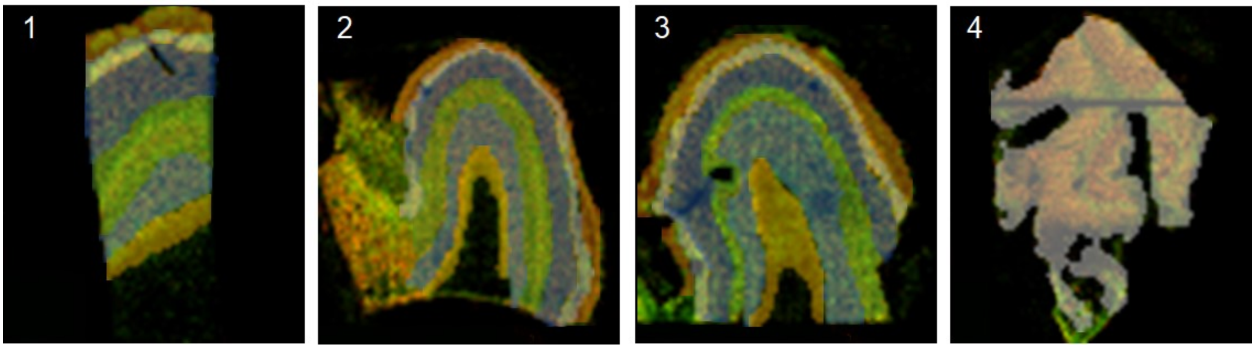


**Supplementary Figure 2**: **Layers' assessment**. Layer masks (L1 red, L2 white, L3 blue, L4 green, L5 light blue, L6 yellow) of the middle plane of samples 1,2, and 3. Grey matter mask (in grey) is shown for sample 4.

#### 3. Convolutional Neural Networks (CNN) parameters

The 2.5D CNN was implemented using the Aliquis® software ecosystem (https://www.bioretics.com/aliquis) version 2.4.3 with TensorFlow backend (https://www.tensorflow.org). Aliquis® is available online free of charge in capped mode.

The **network architecture** is as follows:

```
input 32x32x2 8-bit (promoted to float4 and remapped in range [0 1])
conv (relu) 32 5x5 + max pooling 2x2
conv (relu) 64 3x3 + max pooling 2x2
conv 64 3x3 (relu)
fc (relu) 128 + dropout (0.5)
fc (relu) 128 + dropout (0.5)
fc 2 (softmax)
```

The **network parameters** are as follows:

| Layer name (type) | Output | N. params |
|---|---:|---:|
| conv2d_1 (Conv2D) | 32 | 1632 |
| max_pooling2d_1 (MaxPool 2) | 32 | 0 |
| conv2d_2 (Conv2D) | 64 | 18496 |
| max_pooling2d_2 (MaxPool 2) | 64 | 0 |
| conv2d_3 (Conv2D) | 64 | 36928 |
| conv2d_4 (Conv2D) | 128 | 131200 |
| dropout_1 (Dropout) | 128 | 0 |
| conv2d_5 (Conv2D) | 128 | 16512 |
| dropout_2 (Dropout) | 128 | 0 |
| output (Conv2D) | 2 | 256 |

Total params: 205,024
Trainable params: 205,024
Non-trainable params: 0

**Optimizer hyper-parameters** are as follows:

```
type: SGD (mini-batch stochastic)
batch size: 256
epochs: 300
scale: 0.003921568627 (= 1/255)
learning rate: 0.01
weight decay: 0.00001
momentum: 0.9
loss: infogain_categorical_crossentropy
infogain weight matrix: 0.7; 0; 0 ;1
```

The **datasets** extracted from the raw images are as follows:

Summary **per-image dataset**:
training = 112 (80%); validation = 14 (10%); test = 14 (10%)

Summary **per-patch dataset** (number of 32x32x2 samples):
Total = 2'293'760; background 0 = 2'147'474 (94%); neuron 1 = 146'286 (6%)

Detailed **per-patch dataset** (number of 32x32x2 samples):
Training set: background 0 = 1'717'979; neuron 1 = 117'029
Validation set: background 0 = 214'747; neuron 1 = 14'629

Only the training dataset has been data-augmented.

### 4. CNN statistical assessment and grouping effect

#### 4.1 CNN statistical assessment

The CNN assessment detailed information is listed in the attached file (2.5D assessment.zip). Data are displayed for the four representative stacks of 100 x 100 x 100 $\mu m^3$, both for the manual segmentation (GT) and CNN automatic recognition. Performances are indicated for each neuron.

The manual segmentation has been performed on LAIRA® (https://laira.bioretics.com) and stored in the Ximage open-source format (https://github.com/bioretics/ximage). LAIRA® is available free of charge in trial mode.

#### 4.2 Grouping effect

The 3D reconstruction of the meshes obtained through the 2.5D approach is sometimes characterized by a grouping effect: neurons that are too close to each other are reconstructed as single units, as shown in Supplementary Figure 3.

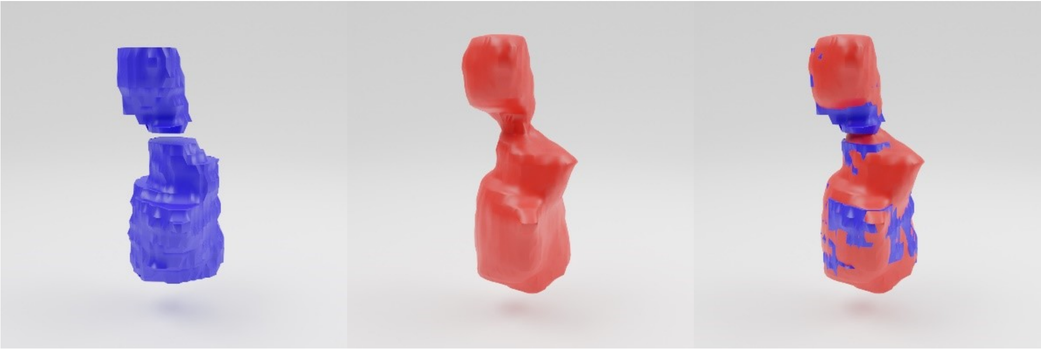

**Supplementary Figure 3: Example of a group.** 3D rendering of two neurons manually segmented (in blue), automatically reconstructed by the 2.5D approach (in red), and the superimposition of the two renderings.

#### 5. Videos legend

**Video1: 2D segmented stack.** Representative stack showing the neuronal segmentation obtained using the CNN. Imaging performed with TPFM, FOV of 450 x 450 $\mu m^2$.In red NeuN neuronal staining in green DAPI nuclei staining.

**Video2: Neuronal 3D reconstruction.** Navigation in the 3D rendering of the meshes of sample 1.
